# Ancient DNA analysis of food remains in human dental calculus from the Edo period, Japan

**DOI:** 10.1101/246868

**Authors:** Rikai Sawafuji, Aiko Saso, Wataru Suda, Masahira Hattori, Shintaroh Ueda

## Abstract

**Objectives:** Although there are many methods for reconstructing diets of the past, detailed taxon identification is still challenging, and most plants hardly remain at a site. In this study, we applied DNA metabarcoding to dental calculus of Edo people for the taxonomic identification of food species.

**Materials and Methods:** DNA was extracted from 13 human dental calculi from the Unko-in site (18th–19th century) of the Edo period, Japan. Polymerase chain reaction (PCR) and sequencing were performed using a primer set specific to the genus *Oryza*, because rice (*Oryza sativa*) was a staple food and this was the only member of this genus present in Japan at that time. DNA metabarcoding targeting plants, animals (meat and fish), and fungi was also carried out to investigate dietary diversity.

**Results:** We detected amplified products of the genus *Oryza* from more than half of the samples using PCR and Sanger sequencing. DNA metabarcoding enabled us to identify taxa of plants and fungi, although taxa of animals were not detected, except human.

**Discussion:** Most of the plant taxonomic groups are present in Japan and include species consumed as food at that time, as confirmed by historical literature. The other groups featured in the lifestyle of Edo people, such as for medicinal purposes and tobacco. The results indicate that plant DNA analysis from calculus provides information about food diversity and lifestyle habits from the past and can complement other analytical methods such as starch analysis and stable isotope analysis.

## 1 INTRODUCTION

Ancient diets have been revealed by multiple methods such as analysis of plant and faunal remains at sites, stable isotope analysis, organic residue analysis of pottery, and morphological analysis of phytoliths and starch grains. Starch grains and phytoliths within ancient calculus (calcified dental plaque) are direct evidence of food items and have revealed dietary habits (Henry, Brooks, & Piperno, 2011; Power, Salazar-García, Straus, González Morales, & Henry, 2015), the spread of domesticated plants (Cristiani, Radini, Edinborough, & Bori, 2016; Madella, García-Granero, Out, Ryan, & Usai, 2014; Piperno & Dillehay, 2008), cooking (Barton & Torrence, 2015), and other usages of teeth (Blatt, Redmond, Cassman, & Sciulli, 2010; Radini et al., 2016). Although the conventional methods are powerful and have been applied for many studies, there are some challenges. For example, taxonomic identification of food at the species or genus level is often difficult, and sometimes the criteria used for assessing this are not completely objective. Moreover, analysis of organs that hardly remain at a site (e.g., leaves, roots, and rhizomes) is almost impossible.

Food DNA analysis of dental calculus has the potential to overcome these limitations. Ancient calculus DNA is one of the richest known sources of ancient biomolecules in the archeological record (Warinner, Speller, & Collins, 2015a). DNA analysis enables detailed taxon identification of plants and animals. In fact, Warinner et al. (2014) and Weyrich et al. (2017) detected plant and animal DNA possibly derived from consumed food, but some challenges with this approach still remain. The efficacy of food DNA analysis of dental calculus has not been adequately validated, and there is a need to improve it as a methodology to analyze the food consumed in the past. Previous studies detected plant and animal DNA from calculus using shotgun sequencing, but the number of individuals was small, and the proportion of plant/animal DNA was quite low. For example, Warinner et al. (2014) reported that DNA within calculus is dominated by bacterial DNA (>99%), with a very small proportion derived from other sources including food DNA. The composition of DNA within calculus was reported to be as follows: 0.002% for animals, 0.005% for fungi, and 0.008% for plants (Warinner et al., 2014). Weyrich et al. (2017) reported that Neanderthal samples contained 0.27% eukaryotic sequences. With such a small proportion, the cost of food DNA analysis is enormous and run-to-run carryover could be a serious problem. Approximately 0.002% carryover contamination (i.e., contamination from previous sequencing runs) was reported using an Illumina sequencer (Bartram et al., 2016; Illumina, 2013). This implies that the risk of misidentification of carryover contamination as food is relatively high because each taxon of food has almost the same proportion of carryover contamination when applying shotgun DNA sequencing to dental calculus.

There is also the matter of databases (Parducci et al., 2017). The level of completeness of reference databases varies by genomic region, which may cause misidentification of taxa. For example, the numbers of species/genera with complete chloroplast genomes were 2,255/1,172 in the National Center for Biotechnology Information (NCBI) RefSeq database (as per the release of 15 September 2017); meanwhile, the approximate numbers of species/genera with depositions of the trnL region of chloroplasts were 72,587/11,037 (downloaded from GenBank on 25 October 2017). Thus, DNA metabarcoding using regions for which abundant information is contained in databases would be more suitable for taxon identification than shotgun sequencing.

In this study, we analyzed food DNA in ancient calculus using DNA metabarcoding analysis, in order to overcome the above-mentioned problems. DNA metabarcoding is a method using a standardized DNA region as a tag for accurate and simultaneous identification of many taxa (Pompanon et al., 2011; Taberlet, Coissac, Pompanon, Brochmann, & Willerslev, 2012). It is often used in the field of ecology for characterizing diet content (De Barba et al., 2014; Shehzad et al., 2012) and has been applied to ancient DNA analysis for taxon identification of bulk bone (Grealy, Douglass, et al., 2016a; Grealy, Macken, et al., 2016b; Murray et al., 2013), permafrost (Willerslev et al., 2014), and lake sediment (Epp et al., 2015; Parducci et al., 2012). By applying this method to calculus samples of Edo people, we investigated whether ancient calculus contains a variety of candidate food DNA including that from staple food.

## 2 MATERIALS AND METHODS

### 2.1 Sample sites

The Unko-in site is the former graveyard of the Unko-in temple at Fukagawa,Tokyo (TAISEI ENGINEERING Co., Ltd., 2010). The excavation was conducted in 1955, and more than 200 skeletons were excavated. In terms of the chronological age of the materials, they were dated to the latter half of the Edo period (from the 18th to the 19th century), as determined from the accompanying cultural finds and historical documents about the temple (Dodo, 1975; Suzuki, Sakura, & Ehara, 1957). The people buried at the temple were townspeople of Edo City, as inferred from their graves and the posthumous Buddhist names (*kaimyo* in Japanese) written on them (Suzuki, 1963). The skeletons are housed at the University Museum, the University of Tokyo (UMUT).

### 2.2 Sampling

Supragingival calculi were collected from the teeth of 14 human adult skeletons using a sterilized dental explorer (YDM Corporation), the identity of which was confirmed by morphological observation (Figure 1). Sampling was performed at UMUT. Calculus from each individual was collected separately into a 1.5 ml DNA LoBind tube (Eppendorf). For comparison, a soil sample from the mandibular foramen was also collected and subjected to the same analysis as the calculus, as a control. Masks, gloves, hairnets, and laboratory coats were worn throughout the process.

**Figure 1:**
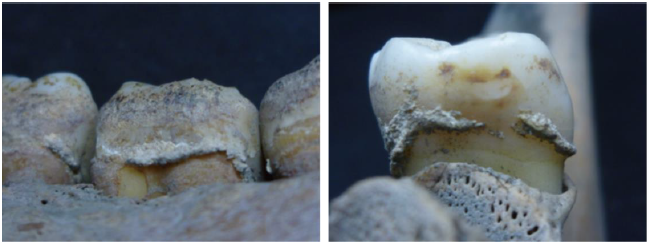
Close-up views of dental calculus on the teeth from the sampled individuals of the Unko-in site.

### 2.3 General equipment

DNA extractions and library preparations were carried out in a laboratory that was physically separated from the one where polymerase chain reaction (PCR) cycling was conducted. Masks, gloves, hairnets, and laboratory coats were worn throughout the process and were replaced regularly. Neither latex gloves nor woolen clothes were used, in order to avoid contamination (Velsko & Warinner, 2017). Items such as tubes and pestles were UV-irradiated before use. Workspaces were wiped with 5% bleach or DNA Away (Thermo Fisher Scientific) and rinsed with 80% ethanol. Laboratory equipment was UV-irradiated, treated with 5% bleach or DNA Away, and rinsed with 80% ethanol. Filtered pipette tips were used in all steps. All DNA extractions and PCR reactions included negative controls.

### 2.4 Confirmation of oral bacteria

To verify the presence of oral bacteria in dental calculus, we performed 16S rRNA analysis. DNA was extracted from one calculus sample from Unko-in site using 1 ml of 0.5 M EDTA and the Fast DNA spin kit for soil (MP Biomedicals), followed by ethanol precipitation. PCR was performed using the primers 27Fmod (5 ′ - AGRGTTTGATYMTGGCTCAG-3 ′) and 338R (5 ′ -TGCTGCCTCCCGTAGGAGT-3 ′) (Kim et al., 2013). The amplification was carried out in 1 × Ex Taq PCR buffer (50 μl), with dNTPs (2.5 mM), Ex Taq polymerase (Takara), each primer (10 μM), and 8.0 μl of template DNA. The cycling profile included an initial denaturation step at 96°C for 2 min; followed by 25 cycles of 96°C for 30 s, 55°C for 45 s, and 72°C for 1 min; and final extension at 72°C for 10 min. Negative controls were included in the PCR amplification. PCR amplicons were purified using the Agencourt AMPure XP kit (Beckman Coulter Genomics), quantified using the Quant-iT PicoGreen dsDNA Assay Kit (Life Technologies), and then sequenced using either 454 GS FLX Titanium or 454 GS JUNIOR (Roche Applied Science). Data analysis was performed as previously described (Said et al., 2014).

### 2.5 DNA extraction

The extraction procedure was a modified version of that reported by Dabney et al. (2013) and Ozga et al. (2016). Dental calculus samples were UV-irradiated for 1 min on each side. After washing twice with molecular-grade water, samples were resuspended for several hours at 55°C in 500 μl of a buffer of 0.5 M EDTA. Samples were then homogenized using each sterile plastic pestle (AS ONE). The digestion step was performed by adding 400 μl of a buffer of 0.5 M EDTA and 10% proteinase K, followed by 8–12 h of digestion at 55°C and 24 h of digestion at 37°C on a rotator. Remaining precipitation was then pelleted by centrifugation in a bench-top centrifuge for 15 min at maximum speed. The supernatant was added to 13 ml of binding buffer, which contained final concentrations of 5 M guanidine hydrochloride, 40% (vol/vol) isopropanol, 0.05% Tween-20, and 90 mM sodium acetate (pH 5.2). A binding apparatus was constructed by forcefully fitting an extension reservoir removed from a Zymo-Spin V column (Zymo Research) into a MinElute silica spin column (Qiagen). The extension reservoir-MinElute assembly was then placed into a 50 ml falcon tube. The 14 ml solution containing binding buffer and the extraction supernatant was then poured into the extension reservoir, and the falcon tube cap was secured. The binding apparatus was centrifuged for 5 min at 1,600 *g*. The extension reservoir-MinElute column was removed from the falcon tube and placed into a clean 2 ml collection tube. The extension reservoir was then removed, and two washing steps were performed by adding 720 μl of binding buffer to the MinElute column, centrifuging on a bench-top centrifuge, and discarding the flow-through. Then, two washing steps were performed by adding 720 μl of PE buffer (Qiagen) to the MinElute column, centrifuging on a bench-top centrifuge, and discarding the flow-through. The MinElute column was dry-spun for 1 min at maximum speed (15,000 rpm) in a bench-top centrifuge and then placed in a fresh 1.5 ml DNA LoBind tube (Eppendorf). For elution, 15 μl of EB buffer was pipetted onto the silica membrane and after 5 min of incubation was collected by centrifugation for 1 min at maximum speed. This step was repeated for a total of 30 μl of DNA extract. One microliter of each extract was quantified using a Qubit High Sensitivity dsDNA assay (Life Technologies).

### 2.6 PCR of the genus *Oryza* (*atpE* gene)

For the detection of rice DNA in calculus samples, PCR was performed using *atpE* gene primers, the sequences of which were specific to the genus *Oryza atpE* gene: atpE_F1 (5′-CGTATTCTCAAGGGACCCATATCT-3′) and atpE_R1 (5′-GCCAAATTGGCGTATTACCAA-3′) (Tozawa et al., 2007). This pair of primers was selected on the basis of the following criteria: (1) The amplified sequence is *Oryza*-specific, as confirmed by BLAST (blastn-megablast) (Altschul, Gish, Miller, Myers, & Lipman, 1990). (2) The amplified sequence is shorter than 100 bp (expected size: 70 bp) because of the highly fragmented nature of ancient DNA (Carpenter et al., 2013). (3) The sequence is present in the chloroplast, mitochondrial, and nuclear genome, because they are usually present in high copy numbers and the sequence is expected to be easily amplified. (4) The primer sequences are largely divergent from sequences of bacteria, archaea, and fungi, to avoid nonspecific amplification.

Each PCR reaction mixture contained 22.6 μl of molecular-grade water, 4 μl of 10 × PCR buffer, 1.0 U AmpliTaq Gold DNA Polymerase (Applied Biosystems),3.2 μl of MgCl_2_ (25 mM), 4 μl of dNTPs (2 mM), 2 μl of each primer (10 μM), and 2.0 μl of DNA template (5 ng/μl) for a total volume of 40 μl. PCR thermal cycling conditions were as follows: 9 min at 95°C; 40 cycles of 30 s at 95°C, 30 s at 50°C, and 30 s at 72°C; and, finally, 7 min at 72°C. Negative controls (water) were included in the PCR amplification, in order to verify the PCR efficiency and to detect contamination, if any. After the PCR amplification, 4–9 μl of PCR solution was loaded on MCE-202 MultiNA (Shimadzu), a capillary microchip electrophoresis system for DNA analysis. For sequencing, second PCR was performed using 1 μl of product from the first PCR with the same conditions as before, but for 10 cycles. Nucleotide sequences of the PCR products were obtained and analyzed in an ABI 3730xl DNA Analyzer (Applied Biosystems) or Applied Biosystems 3130xl Genetic Analyzer by the Fasmac sequencing service (Fasmac). Sequencing was conducted in both directions.

### 2.7 DNA metabarcoding

DNA extracts and extraction blank controls were amplified using four DNA metabarcoding markers: *trn*L intron (P6 loop of the chloroplast *trn*L intron) for plants, 12S rRNA (two different primer sets) for vertebrates/teleosts, and ITS1 for fungi. As for animal DNA metabarcoding, it is expected that human DNA would be amplified predominantly when targeting animal DNA, so we used human blocking primer sets for DNA metabarcoding with 12S rRNA primer sets. The sequences of primers used for DNA metabarcoding are listed in Table 2.

**Table 2:**
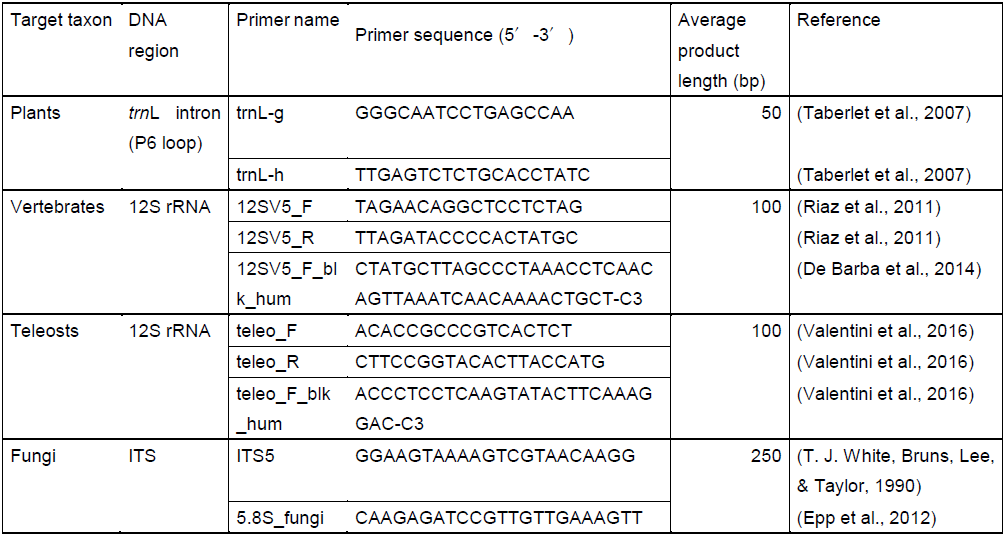
List of primers used for DNA metabarcoding in this study

All primers were modified to include a subset of Nextera XT (Illumina) adapters. Each PCR reaction mixture of the first PCR contained 11.38 μl of molecular-grade water, 2.5 μl of 10 × PCR buffer, 0.6 U AmpliTaq Gold DNA Polymerase (Applied Biosystems), 2 μl of MgCl_2_ (25 mM), 2.5 μl of dNTPs (2 mM), 1.25 μl of each primer (10 μM), and 4 μl of DNA template (5 ng/μl) for a total volume of 25 μl. PCR thermal cycling conditions were as follows: 9 min at 95°C; 40 cycles of 30 s at 95°C, 30 s at 50°C, and 30 s at 72°C; and finally 7 min at 72°C. PCR products were purifide using Agencourt AMPure XP kit (Beckman Coulter Genomics) and quantified using the HS dsDNA Qubit Assay on a Qubit 2.0 Fluorometer (Life Technologies).

The second-round PCR (second PCR) used the first PCR products as a template. Each PCR reaction mixture of the first PCR contained 25 μl of 2 × KAPA HiFi HotStart ReadyMix (KAPA Biosystems), 5 μl of each Nextera XT Index Primer 1, 5 μl of each Nextera XT Index Primer 2, and 15 μl of the first PCR products as a template, for a total volume of 50 μl. PCR products were purified using Agencourt AMPure XP kit (Beckman Coulter Genomics), quantified using the HS dsDNA Qubit Assay on a Qubit 2.0 Fluorometer (Life Technologies), and visualized using a High Sensitivity DNA Assay Chip kit on a Bioanalyzer 2100 (Agilent). Samples with less than 0.5 ng/μl DNA were discarded.

Purified PCR products were pooled to equimolar concentration (4 nM). Five microliters of the pool library was denatured with 5 μl of fresh 0.1 N NaOH. Including HT1 buffer (provided by Illumina), the denatured library was diluted to a final concentration of 4–8 pM. Here 5% PhiX DNA spike-in control was added to improve data quality of low-diversity samples such as PCR amplicons (Carpenter et al., 2013; Miya et al., 2015). The libraries of paired-end reads were sequenced with Illumina MiSeq (Illumina Inc.) using MiSeq Reagent Kit Nano and Micro v2 (2 × 150 bp).

### 2.8 Data analysis

Nextera XT adapters were removed from paired-end reads using cutadapt v.1.11 (Martin, 2011). Trimmed paired-end reads were then merged using the *illuminapairedend* tool in OBITOOLS (Boyer et al., 2016). Unmerged reads were removed using *obigrep*. Sequences with counts ≤10 were removed using *obiuniq, obistat*, and *obigrep*. Each sequence was then assigned the status of “head,” “internal,” or “singleton” using *obiclean*. Since all sequences labeled “internal” probably correspond to PCR substitutions and indel errors, only “head” and “singleton” sequences were used for sequential taxonomic assignment. All primer sequences were removed because mutations may be inserted in the process of PCR amplification. Taxonomic assignments were identified using blastn megablast on the NCBI website (https://blast.ncbi.nlm.nih.gov/Blast.cgi) with the database of the NCBI nucleotide collection (nr/nt) (Altschul et al., 1990). For plant (*trn*L primer sets) and animal (12S rRNA primer sets) DNA metabarcoding, taxa with 100% query coverage and 100% identity were accepted. For fungal (ITS primer sets) DNA metabarcoding, taxa with 99% identity were accepted. Identification was determined from the genus to the order level from accepted taxa.

The taxa detected from soil samples were removed from among those used for the analysis of calculus samples. Functional annotations of fungal genera were determined by the descriptions of Tedersoo et al. (2014).

## 3 RESULTS

### 3.1 DNA extraction

We extracted DNA successfully from 13 samples out of 14 samples. DNA extraction yields are shown in Table 1. The total amount of DNA was 206– 1,650 ng and normalized yields of DNA were 13–90 ng/mg calculus, which is far more than for DNA extracted from bone and dentine (Warinner, Speller, Collins, & Lewis, 2015b). We also confirmed the presence of oral bacteria such as *Streptococcus parasanguinis* and *Streptococcus salivarius* (Supporting Information Table S1).

**Table 1:** Ancient dental calculus information and DNA extraction yields

| Individuals | Sex | Sample<br>(mg) | Total DNA (ng) | Normalized DNA<br>yield (ng/mg) |
| --- | --- | --- | --- | --- |
| wn2016-F01 | Female | 11 | 345 | 31.4 |
| wn2016-F04 | Female | 16 | 206 | 12.9 |
| wn2016-F07 | Male | 20 | 627 | 31.4 |
| wn2016-F08 | Female | 16 | 1434 | 89.6 |
| wn2016-F10 | Male | 29 | 690 | 23.8 |
| wn2016-F12 | Female | 35 | 1410 | 40.3 |
| wn2016-F23 | Male | 19 | 333 | 17.5 |
| wn2016-F24 | Male | 17 | 699 | 41.1 |
| wn2016-F37 | Female | 29 | 1476 | 50.9 |
| wn2016-F39 | Male | 33 | 930 | 28.2 |
| wn2016-F41 | Male | 43 | 1650 | 38.4 |
| wn2016-F43 | Female | 23 | 1050 | 45.7 |
| wn2016-F44 | Female | 12 | 1038 | 86.5 |

### 3.2 Plants

We examined whether DNA of the genus *Oryza*, which includes rice species (*Oryza sativa*), was detected from ancient calculus in Edo people. This is because rice was a staple food of people living in Edo City and is likely to be detected from such calculus. DNA amplification of *Oryza* was detected in eight out of 13 calculus samples by PCR, as shown in Figure 2. The sequences of the PCR products were identified as the genus *Oryza* (E value = 2.0 × 10^−26^), which included a cultivated rice taxon (*O. sativa*). There was no significant difference between the sexes (Fisher’s exact test, *p* = 0.59).

**Figure 2:**
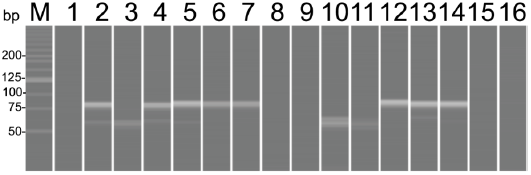
PCR amplification products of dental calculus using *Oryza atpE* gene primer sets. M, molecular weight markers; 1, wn2016-F01; 2, wn2016-F04; 3, wn2016-F07; 4, wn2016-F08; 5, wn2016-F10; 6, wn2016-F12; 7, wn2016-F23; 8,negative control; 9, wn2016-F24; 10, wn2016-F37; 11, wn2016-F39; 12, wn2016-F41; 13, wn2016-F43; 14, wn2016-F44; 15, soil from wn2016-F39; 16, negative control.

Next, we investigated whether other genera of plants could be detected from ancient calculus samples using DNA metabarcoding analysis (shown in Table 3). For DNA metabarcoding, the P6 loop of the chloroplast *trn*L intron was amplified using the primers trnL-g and trnL-h (Taberlet et al., 2007), as shown in Table 2. After identification with Blast, seven taxa were confirmed at the family level and 10 taxa were confirmed at the genus level from six samples in total, as shown in Table 4. Sequences of the family Fagaceae were detected from four individuals, and sequences of the family Poaceae and the genera *Lactuca, Celtis*, and *Oryza* were detected from two individuals. Other taxa were detected from one individual. The sequence of *Ginkgo biloba* was detected from both soil and calculus DNA, and we excluded this sequence from further analyses.

**Table 3:** Results of DNA metabarcoding

| Individuals | Plants<br>(trnL) | Vertebrates<br>(12S rRNA) | Teleosts<br>(12S rRNA) | Fungi<br>(ITS) |
| --- | --- | --- | --- | --- |
| wn2016-F04 | ✓✓ | - | - | * |
| wn2016-F08 | - | *Homo* | - | ✓✓ |
| wn2016-F10 | ✓✓ | - | - | ✓✓ |
| wn2016-F12 | ✓✓ | - | - | ✓✓ |
| wn2016-F23 | - | - | - | - |
| wn2016-F41 | ✓✓ | - | - | - |
| wn2016-F43 | ✓✓ | - | - | ✓✓ |
| wn2016-F44 | ✓✓ | - | - | ✓✓ |
| Soil | ✓✓ | - | - | ✓✓ |

Experiments that did not produce any identified sequences are shown by a hyphen. An asterisk indicates that the experiment was not performed. In the wn2016-F08 sample, human DNA was amplified with the primer set for vertebrates, which is represented as *Homo*. The soil sample was obtained from mandibular foramen of wn2016-F39.

**Table 4:**
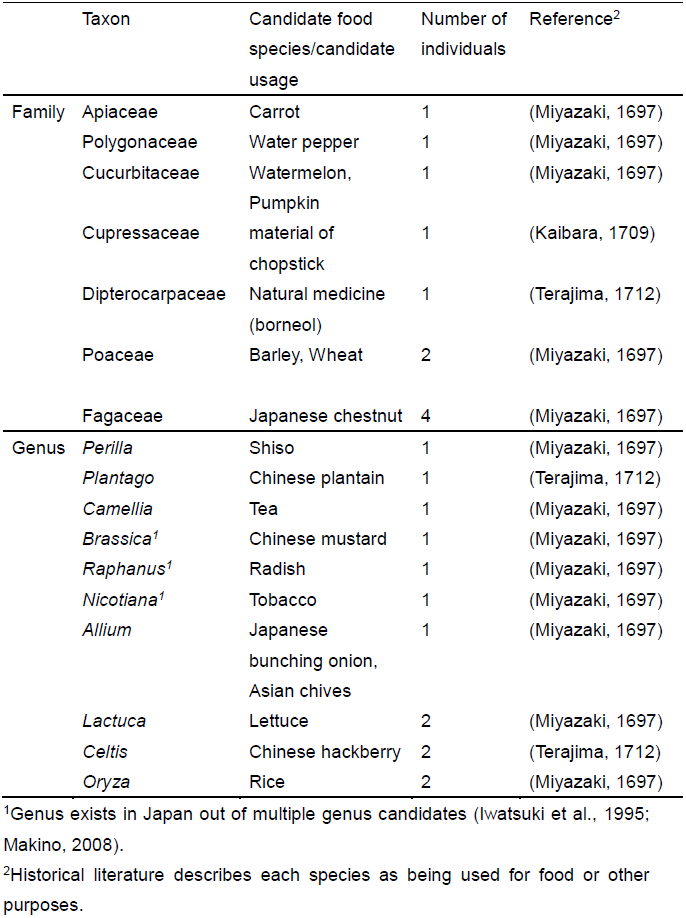
Plant taxa identified using *trn*L primer set

### 3.3 Animals

We also investigated whether animal DNA could be detected from ancient calculus samples. We used “12SV5” primer pairs for the amplification of vertebrate DNA and “teleo” primer pairs for the amplification of teleosts (Riaz et al., 2011; Valentini et al., 2016). The results are shown in Table 3. No animal taxon except human was detected from the calculus samples. Human DNA was detected from one sample.

### 3.4 Fungi

With regard to fungi, 3 taxa were confirmed at the order level, 4 taxa at the family level, and 12 at the genus level, from five calculus samples in total, as shown in Table 5. Sequences of the genus *Wallemia* were detected from four individuals, and sequences of the order Filobasidiales and the genera *Cladosporium, Peniophora, Sporidiobolus*, and *Trametes* were detected from two individuals. Other taxa were detected from one individual. Sequences of the genera *Aspergillus* and *Malassezia* were detected from both soil and calculus DNA, and we excluded these sequences from further analyses.

**Table 5:** Fungal taxa identified using ITS primer set

|  | Taxon | Trophic status | Functional group | Number of individuals |
| --- | --- | --- | --- | --- |
| Order | Agaricales |  |  | 1 |
|  | Hypocreales |  |  | 1 |
|  | Filobasidiales |  |  | 2 |
| Family | Coriolaceae |  |  | 1 |
|  | Diatrypaceae |  |  | 1 |
|  | Didymellaceae |  |  | 1 |
|  | Sporidiobolaceae |  |  | 1 |
| Genus | *Alternaria* | Biotroph | Plant pathogen | 1 |
|  | *Clonostachys* | Biotroph | Plant pathogen | 1 |
|  | *Perenniporia* | Saprotroph | White rot | 1 |
|  | *Phaeosphaeria* | Saprotroph |  | 1 |
|  | *Phanerochaete* | Saprotroph | White rot | 1 |
|  | *Porostereum* | Saprotroph | White rot | 1 |
|  | *Rhodotorula* | Saprotroph |  | 1 |
|  | *Cladosporium* | Saprotroph |  | 2 |
|  | *Peniophora* | Saprotroph | White rot | 2 |
|  | *Sporidiobolus* | Biotroph | Mycoparasite | 2 |
|  | *Trametes* | Saprotroph | White rot | 2 |
|  | *Wallemia* | Saprotroph |  | 4 |

## 4 DISCUSSION

The purpose of this study was to assess the potential utility of ancient calculus DNA as a source of dietary evidence. Indeed, we demonstrated that plant DNA can be extracted from ancient calculus and identified at the family or genus level. Most of the identified taxa included species that were described as food in the historical literature of that time (Table 4). For example, lettuce (*Lactuca sativa*) was described as the food plant *chisha* (in Japanese) in *Nogyo Zensho,* published in the Edo period (Miyazaki, 1697).

In particular, we detected *Oryza* sequences by both genus-specific PCR and DNA metabarcoding. Amplification was observed from more than half of the calculus samples in genus-specific PCR, but not from soil samples. Wild *Oryza* species are not distributed in Japan, so these results suggest that the *Oryza* DNA detected from calculus samples is derived from rice consumed as food.

The diet of townspeople in Edo City has been investigated by analyzing historical literature and performing stable isotope analysis. Historical studies suggested that the diet of the Edo townspeople was mainly composed of rice and vegetables, accompanied by fish (Ayako Ehara, Ishikawa, & Syoko, 2009). Stable isotope analysis revealed that C3-based terrestrial food (e.g., rice and vegetables), freshwater fish, and marine fish were their main sources of dietary protein (Tsutaya, Nagaoka, Kakinuma, Kondo, & Yoneda, 2016). It is consistent with these reports that DNA of the genus *Oryza* and taxa including a variety of vegetables was detected from calculus samples in this study, although fish DNA was not detected. DNA metabarcoding of ancient calculus samples thus appears to be a promising approach for screening the diversity of plant food from the past.

### 4.1 Plants

For the genus *Oryza*, there is a difference between the results of PCR obtained with the *Oryza*-specific primer set and DNA metabarcoding with the *trn*L primer set. PCR products were detected in more than half of the samples (eight out of 13 calculus samples) when using the specific primer set targeting sequences on mitochondria, chloroplasts, and the nucleus. On the other hand, a sequence of the genus *Oryza* was detected from only two out of eight samples when using the chloroplast *trn*L primer set for DNA metabarcoding analysis. This can be explained by the number of genome regions in which the sequences exist. The numbers of mitochondrial and chloroplast DNA copies contained in each organelle range from 10 to more than 100. Moreover, multiple copies of both organelles are contained in a cell, so the sequences present in both mitochondrial and chloroplast genomes are very easy to amplify by PCR. We selected this specific primer set because the sequences exist in multiple regions, including two sites of the mitochondrial genome, one site of the chloroplast genome, and eight sites of the nuclear genome. This can explain why the sequences of the genus *Oryza* were detected in the experiment with the specific primers much more than in that with the *trn*L primer set.

DNA of the genus *Allium* was detected in this study. Darbyshire & Henry (1981) reported that no starch was detected from *Allium* species, although fructans were present. Shibutani (2015) also reported that many Japanese *Allium* species produced a relatively small number of starch grains, so it seems to be quite difficult to identify *Allium* species by starch grain analysis of archeological materials. Our results indicate that plant DNA analysis from calculus enables us to identify even plants hardly remain at a site or plants that produce little starch.

Taxa that are difficult to interpret as food were also detected from the calculus samples (Table 4). Wild species of the genus *Nicotiana* did not exist in Japan (Iwatsuki, Boufford, & Ohba, 1995), and only cultivated species for making tobacco were present at that time. Smoking was common among the townspeople in Edo City (Kitagawa, 1853), so it seems reasonable that this taxon was detected from the calculus samples, although a pipe was normally used and chewing tobacco has not been recorded.

The source of the plant of the Cupressaceae family is ambiguous. Hardy et al. (2017) detected plant fibers from a hominin (1.2 million years ago) dental calculus adjacent to an interproximal groove, indicative of oral hygiene activity such as tooth picking. The sequence of the Cupressaceae family could also have been derived from chopsticks or tooth picking, although this is unclear.

The detection of plants of the Dipterocarpaceae family is surprising because these trees are well-known constituents of Asian rainforests (Dayanandan, Ashton, Williams, & Primack, 1999) and do not inhabit Japan. Therefore, this cannot be explained without the existence of trade. One plausible explanation is that this was derived from impurities of borneol (*ryunou* in Japanese), chemical compounds extracted from *Dryobalanops aromatica* (*ryunou-jyu* in Japanese), belonging to the family Dipterocarpaceae. Borneol is one of the traditional herbal medicines and was commonly used as a component of tooth powder in the Edo period. Tooth powder in Edo City was made mainly from *bousyu-zuna* (fine sand from Chiba Prefecture), flavored with borneol and clove (Kitamura, 1830). Toothbrushing was common among the townspeople of Edo City, so DNA of Dipterocarpaceae could have been derived from borneol in tooth powder.

Detecting evidence of ancient oral hygiene activities is often difficult. It has been considered that interproximal wear grooves on fossil teeth could have been caused by tooth picking in order to extract food stuck between teeth, though a number of other hypotheses for the cause of the grooves have been proposed (Lozano, Subirà, Aparicio, Lorenzo, & Gómez-Merino, 2013; Ungar, Grine, Teaford, & Pérez-Pérez, 2001). Previous studies analyzed microfossils and chemical compounds from dental calculus and suggested the performance of oral care or the use of a toothbrush or toothpick (Buckley, Usai, Jakob, Radini, & Hardy, 2014; Cummings, Yost, & Sołtysiak, 2016). We found DNA that could be derived from tooth powder from ancient dental calculus, though more research is necessary to determine the origin of the Dipterocarpaceae DNA.

### 4.2 Animals

Analysis of faunal food residues by DNA metabarcoding requires one more step. There is a problem that samples such as feces and calculus are often enriched with predator DNA (i.e., humans in this study), so the result of PCR amplification could be dominated by predator DNA rather than by prey DNA (Piñol, Mir, Gomez-Polo, & Agustí, 2015). To overcome this problem, a “blocking primer” has been used to reduce the amplification of predator DNA. This primer preferentially binds to predator DNA and is modified so that it does not prime amplification. There are various types of blocking primer, and the most effective and common type is the primer that overlaps with the 3′ end of the universal primer but extends into predator-specific sequence and is modified with a C3 spacer (three hydrocarbons) at the 3′ end (Vestheim & Jarman, 2008). This C3 spacer prevents elongation during PCR, so prey DNA is preferentially amplified.

In this study, we used a blocking primer of human DNA, but meat or fish DNA could not be identified from the calculus samples. Previous studies reported that blocking primer inhibits the amplification of target DNA in some cases (Piñol et al., 2015; Port et al., 2016). We reduced the concentration of human blocking primers considering those studies. However, there is a possibility that blocking primers worked as an inhibitor of the PCR reaction because no animal DNA was detected. This is troublesome because human DNA would be amplified without blocking primers. In fact, human DNA was amplified even with blocking primers (Table 3). This dilemma is difficult to resolve, and study using modern calculus is needed to confirm this interpretation and improve the method.

### 4.3 Fungi

Fungal DNA was also detected from the calculus samples (Table 5). It is interesting that the functional group of many taxa was white rot (wood-decaying fungi). The proportion of white-rot fungi in soil is quite small (<1%) (Tedersoo et al., 2014), and we excluded taxa that were detected from soil samples. The source of the detected fungi is unclear, but one possibility is that they came from wood used as chopsticks or a toothbrush. Other identified taxa such as *Alternaria* and *Clonostachys* are plant pathogens, which might have been derived from plants used as food (Weyrich et al., 2017).

### 4.4 Advantages and limitations of this method

The advantages of food DNA analysis of calculus include the possibility of performing genus/species-level identification. Species-level identification was not performed in this study, but it would theoretically be possible if specific primers were used. This method also enables us to detect plant species that hardly remain at a site (e.g., leaves, roots, and rhizomes) and can complement other methods such as stable isotope and starch analyses.

Food DNA in calculus can directly reveal the diversity of food consumed in the past, including in prehistoric times. This analysis can also be used to investigate the existence of trade. In this study, evidence of trade was presented from the identification of plants of the family Dipterocarpaceae. A previous study pointed out that materials within calculus include not only food but also nonfood items relevant to oral hygiene practices (Radini, Nikita, Buckley, Copeland, & Hardy, 2017). For example, the use of medicinal herbs has been discussed in studies of dental calculus using DNA, starch grain, and chemical analyses (Buckley et al., 2014; Hardy et al., 2012; Weyrich et al., 2017). We found DNA that could be derived from tooth powder from an archeological material for the first time. Combining multiple methods such as proteomics (Sawafuji et al., 2017; Warinner et al., 2014), stable isotope analysis (Santana-Sagredo, Lee-Thorp, Schulting, & Uribe, 2014; Tsutaya, 2017), and starch grain analysis (Hardy et al., 2009; Zhang et al., 2017) should lead to a deeper understanding of various facets of human life in the past.

With regard to the merits of DNA metabarcoding, it is cost-effective and suited to population analysis. It is also compatible with screening for food DNA. In the field of sedimentary ancient DNA (*sed*aDNA), DNA metabarcoding is a common method for analyzing vegetation and fauna (Alsos et al., 2016; Pansu et al., 2015), and its use may also spread to ancient calculus studies for dietary analysis

There is a possibility that the nuclear genes or genome can be analyzed from the food debris of calculus. It was reported that high-throughput sequencing technologies are not suitable for analyzing the genome from charred archeobotanicals (Nistelberger, Smith, Wales, Star, & Boessenkool, 2016). Calculus is not charred, so it seems likely that nuclear sequences could be obtained from it. If a taxon of interest is detected by DNA metabarcoding, one can expand the analysis to more specific research, for example, examining nuclear genes or correlations with oral microbiota.

## ACKNOWLEDGMENTS

The author contributions were as follows: R.S. conceived the study, designed the study, collected samples, extracted DNA from the samples, performed the experiments, analyzed the data, and drafted the manuscript; A.S. contributed to collecting samples and estimated the sex; W.S. performed the experiments on oral bacteria and analyzed the data; M.H. provided resources and contributed to analyzing oral bacteria; and S.U. contributed to the design of the study and to drafting the manuscript. All authors gave final approval for publication of this manuscript.

We are grateful to Gen Suwa for allowing us to collect human dental calculus samples in UMUT. Fuzuki Mizuno and Masahiko Kumagai gave us helpful advice on the experimental design for ancient DNA analysis. We thank Ryutaro Jo for providing us with information and documents about the DNA extraction methods from calculus. We also thank Hirokazu Tsukaya for helpful comments on taxon identification and the exploitation of plants. Takumi Tsutaya gave us insightful comments on our research and the manuscript. This work was supported by a Grant-in-Aid from The Manabu Yoshida Memorial Fund for Scientific Studies on Cultural Properties, Tokyo, Japan.

